## Supplementary Figures for "The glycosyltransferase POGLUT1 regulates muscle stem cell development and maintenance in mice"

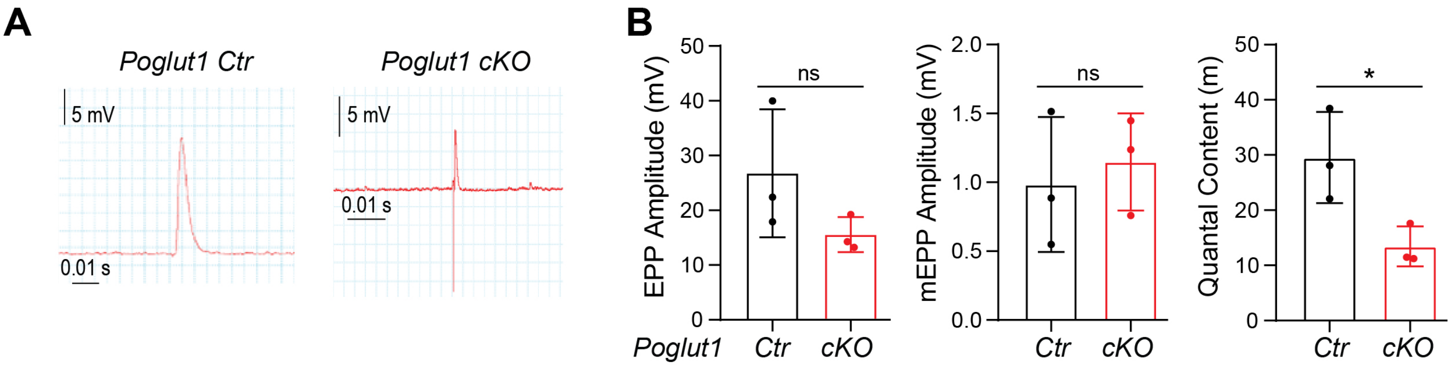


**Supplementary Figure 1. *Poglut1-cKO* muscles exhibit reduced quantal content.**

**A)** Electrophysiological recordings from LAL muscles at low frequency stimulation (0.5 Hz, 100 s).

**B)** Quantification of the EPP amplitude, miniature EPP (mEPP) amplitude, and the quantal content from control and *Poglut1-cKO* muscles. The mean mEPPs amplitude showed no significant difference between control and cKO fibers, suggesting comparable spontaneous release. While the mean size of the EPPs showed no significant difference between cKO and control, the quantal content values of cKO were significantly lower than that in control, suggesting an alteration in neurotransmission in cKO mice (*Poglut1-Ctr*: 3 mice (52 terminals); *Poglut1-cKO*: 3 mice (33 terminals)). In B and D, each dot represents an animal. Mean ± SD is shown. Unpaired *t* test was used. ns: not significant, * *P*<0.05.

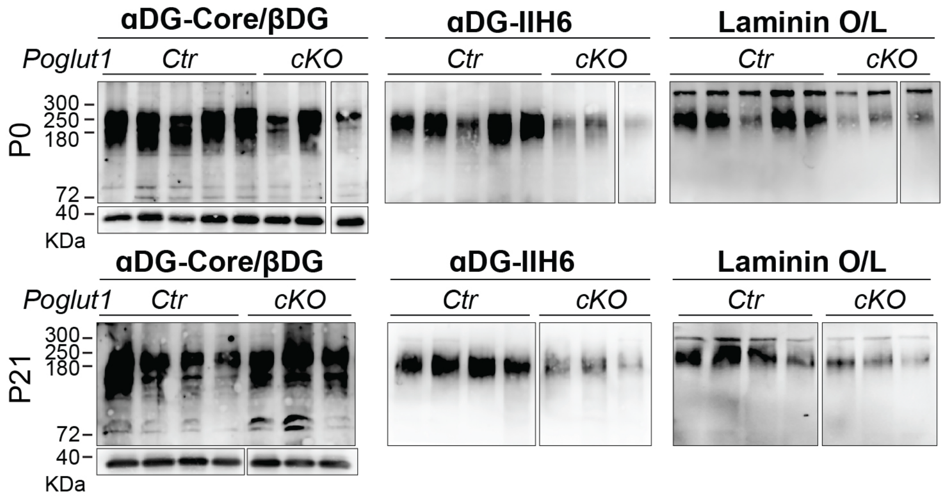


**Supplementary Figure 2.** ***Poglut1-cKO*** **muscles exhibit reduced αDG glycosylation and laminin binding.** Shown is α-DG glycosylation status in *Poglut1-cKO* and control mouse muscles at P0 and P20 assessed by western blotting using antibodies against the glycosylated form of α-DG (αDG-IIH6) and the core α-DG, and by laminin overlay assay. An antibody against β-DG was used to assess loading. Wheat germ agglutinin-enriched muscle lysates from control and cKO mice were used. The western blot showed a reduced expression level of glycosylated α-dystroglycan in cKO mice at P0 and P20, as well as a diminished binding activity in the laminin overlay assay when compared to control mice. Each lane represents an individual animal. Boxes mark samples taken from non-adjacent wells of the same gel.

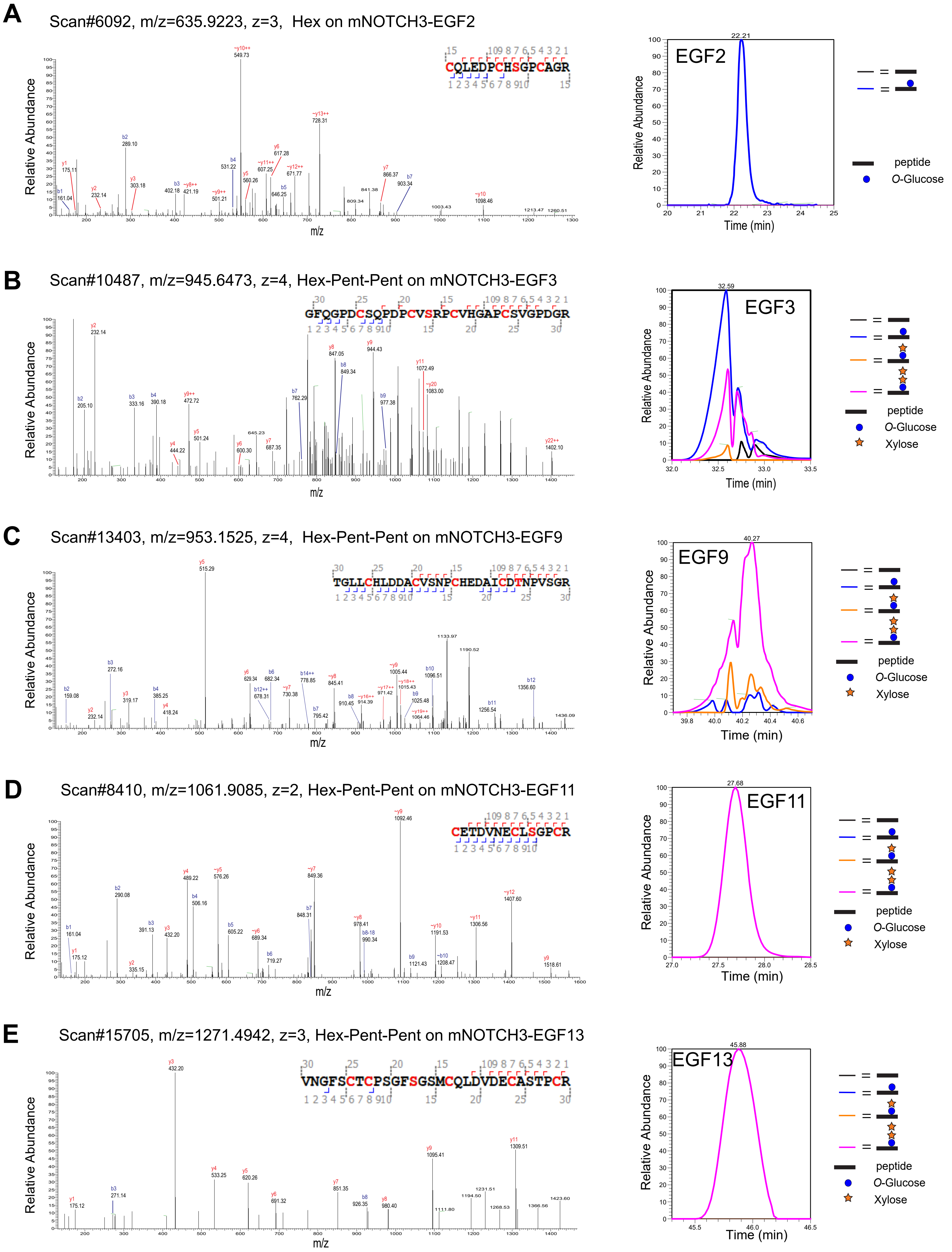
A
Scan#6092, m/z=635.9223, z=3, Hex on mNOTCH3-EGF2
CQLEDPCHSGPCAGR
EGF2
B
Scan#10487, m/z=945.6473, z=4, Hex-Pent-Pent on mNOTCH3-EGF3
GFQGPDCSQPDPCVSRPCVHGAPCSVGPDGR
EGF3
C
Scan#13403, m/z=953.1525, z=4, Hex-Pent-Pent on mNOTCH3-EGF9
TGLLCHLDDACVSNPCHEDAICDTNPVSGR
EGF9
D
Scan#8410, m/z=1061.9085, z=2, Hex-Pent-Pent on mNOTCH3-EGF11
CETDVNECLSGPCR
EGF11
E
Scan#15705, m/z=1271.4942, z=3, Hex-Pent-Pent on mNOTCH3-EGF13
VNGFSCTCPSGFSGSMCQLDVDECASTPCR
EGF13
Relative Abundance
m/z
Time (min)
peptide
O-Glucose
Xylose

**F** Scan#12301, m/z=1230.4945, z=3, Hex-Pent-Pent and Fucose on mNOTCH3-EGF19

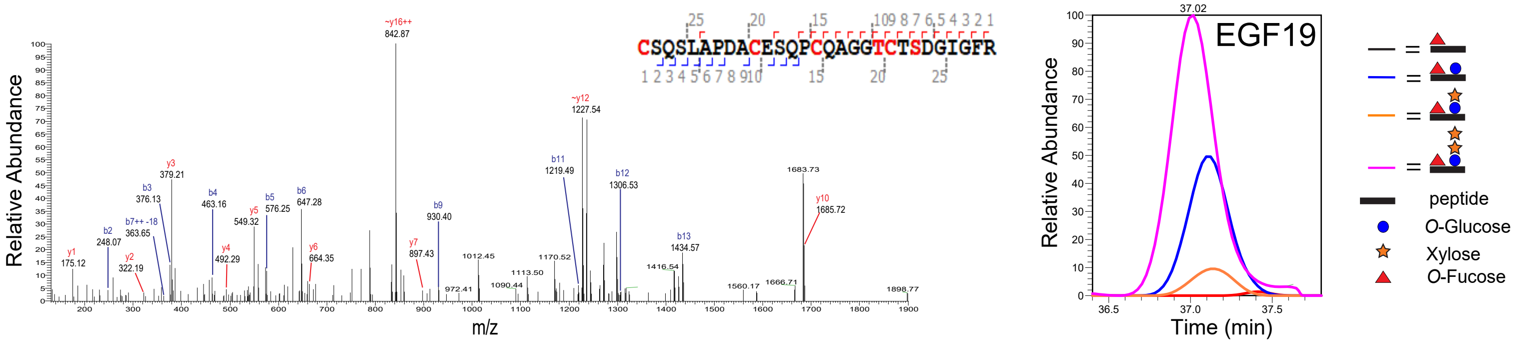


**G** Scan#13640, m/z=1018.4184, z=4, Hex-Pent-Pent and Fucose on mNOTCH3-EGF23

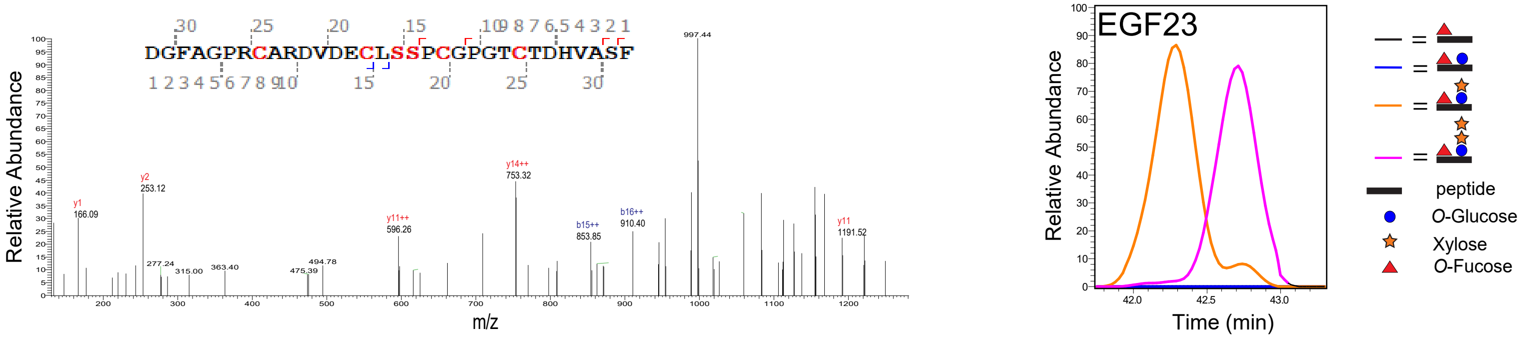


**H** Scan#16575, m/z=1334.5195, z=4, Hex-Pent-Pent and Fucose on mNOTCH3-EGF29

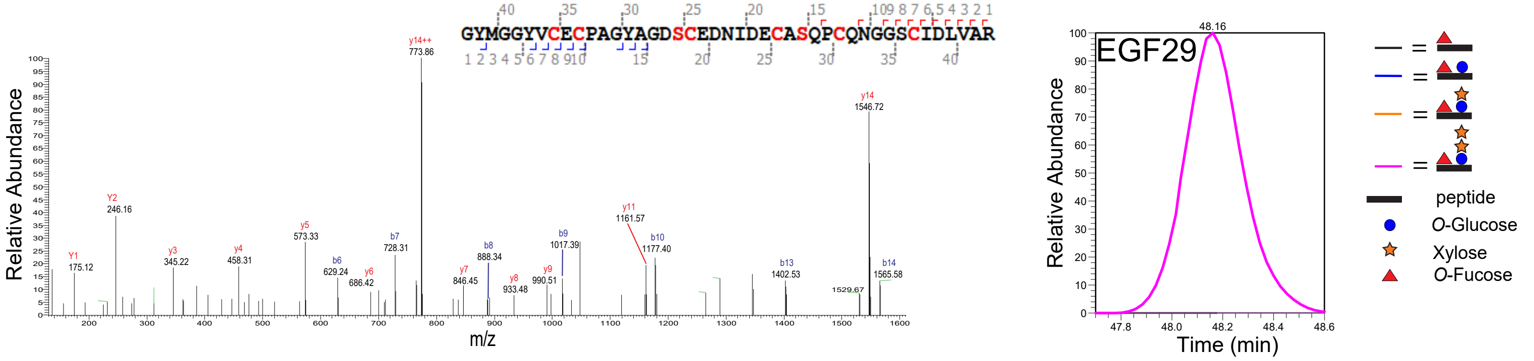


**I** Scan#8273, m/z=933.7157, z=3, Hex-Pent-Pent on mNOTCH3-EGF32

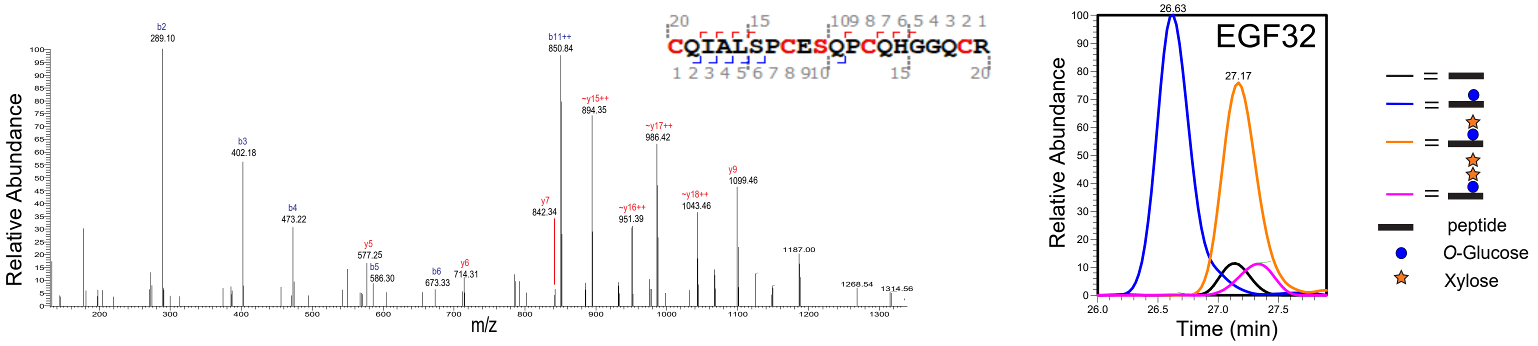

**Supplementary Figure 3. Mass Spectra and Extracted Ion Chromatograms of peptides modified by POGLUT1 from NOTCH3.** Representative MS/MS spectra are shown on the left, and Extracted Ion Chromatograms (EICs) on the right for peptides containing the POGLUT1 consensus site from mouse NOTCH3 EGF repeats. **A)** Peptide with *O*-glucose monosaccharide on EGF2. **B)** Peptide with *O*-glucose-xylose-xylose trisaccharide on EGF3. **C)** Peptide with *O*-glucose-xylose-xylose trisaccharide on EGF9. **D)** Peptide with *O*-glucose-xylose-xylose trisaccharide on EGF11. **E)** Peptide with *O*-glucose-xylose-xylose trisaccharide on EGF13. **F)** Peptide with *O*-glucose-xylose-xylose trisaccharide and *O*-fucose monosaccharide on EGF19. **G)** Peptide with *O*-glucose-xylose-xylose trisaccharide and *O*-fucose monosaccharide on EGF23. **H)** Peptide with *O*-xylose-xylose trisaccharide and *O*-fucose monosaccharide on EGF29. **I)** Peptide with *O*-glucose-xylose-xylose trisaccharide on EGF32. The b- and y-ions are indicated in blue and red font, respectively. ~ indicates ions that lost glucose in the gas phase during fragmentation. Due to the lability of *O*-glycans in the gas phase during collision, these modifications can fall off which can lead to incorrect annotation of the modified amino acid. However, we can predict the correct position based on the putative consensus sequence for modification by POGLUT1.
